# Towards a molecular mechanism underlying mitochondrial protein import through the TOM and TIM23 complexes

**DOI:** 10.1101/2021.08.30.458282

**Authors:** Holly C. Ford, William J. Allen, Gonçalo C. Pereira, Xia Liu, Mark S. Dillingham, Ian Collinson

**Author notes:** MRC - Mitochondrial Biology Unit, University of Cambridge, The Keith Peters Building, Cambridge Biomedical Campus, Cambridge CB2 0XY, UK.

## Abstract

Nearly all mitochondrial proteins need to be targeted for import from the cytosol. For the majority, the first port of call is the translocase of the outer membrane (TOM complex), followed by a procession of alternative molecular machines, conducting transport to their final destination. The pre-sequence translocase of the inner-membrane (TIM23-complex) imports proteins with cleavable pre-sequences, and comes in two flavours: the TIM23^SORT^ complex mediates inner mitochondrial membrane (IMM) protein insertion and the TIM23^MOTOR^ complex delivers proteins to the matrix. Progress in understanding these transport mechanisms has been hampered by the poor sensitivity and time-resolution of import assays. However, with the development of an assay based on split NanoLuc luciferase, we can now explore this process in greater detail. Here, we apply this new methodology to understand how Δψ and ATP hydrolysis, the two main driving forces for import through the TIM23^MOTOR^ complex, contribute to the import of pre-sequence-containing precursors (PCPs) with varying properties. Notably, we found that two major rate-limiting steps define PCP import time: passage of PCP across the OMM and initiation of IMM transport by the pre-sequence. The rates of these steps are influenced by PCP properties such as size and net charge, but not total amount of PCP imported – emphasising the importance of collecting rapid kinetic data, achieved here, to elucidate mechanistic detail. The apparent distinction between transport through the two membranes (passage through TOM is substantially complete before PCP-TIM engagement) is in contrast with the current view that import occurs through TOM and TIM in a single continuous step. Our results also indicate that PCPs spend very little time in the TIM23 channel – presumably rapid success or failure of import is critical for maintaining mitochondrial fitness.

## INTRODUCTION

Mitochondria are double membrane-bound eukaryotic organelles responsible for the biosynthesis of ATP among many other essential cellular functions (Nowinski et al., 2018)(Rouault, 2012)(Nicholls, 1978)(Chen et al., 2003)(Nishikawa et al., 2000)(Hoth et al., 1997)(Chandel, 2015)(Wang and Youle, 2009). Of more than a thousand proteins that constitute the mitochondrial proteome, all but a handful (encoded on the mitochondrial genome - 13 in human) are synthesised in the cytosol, and must be imported. Almost all mitochondrial proteins (exceptions include precursors of α-helical outer mitochondrial membrane (OMM) proteins) initially enter mitochondria *via* the translocase of the outer membrane (TOM complex) which contains the pore-forming β-barrel protein, Tom40 (Ahting et al., 2001)(Guan et al., 2021)(Araiso et al., 2019). From here they are sorted to a number of bespoke protein import machineries, which direct them to their final sub-mitochondrial destination: the OMM, inter-membrane space (IMS), inner-membrane (IMM), or matrix.

Roughly 60-70% of mitochondrial precursor proteins – almost all those targeted to the matrix and a subset of IMM proteins – have a positively-charged, amphipathic α-helical pre-sequence, also known as a mitochondrial targeting sequence (MTS) (Araiso et al., 2019)(Vögtle et al., 2009). These pre-sequence-containing precursors (PCPs) are transferred to the translocase of the inner membrane (TIM23-complex) once their N-termini emerge from the Tom40 channel, and pass through in an unfolded state (Eilers and Schatz, 1986)(Matouschek et al., 1997)(Neupert and Brunner, 2002)(Rassow et al., 1990) (Neupert and Herrmann, 2007). Genetic and biochemical experiments have elucidated the key constituents of the TIM23-complex (Blom et al., 1993) (Maarse et al., 1992) (Emtage and Jensen, 1993) (Maarsea et al., 1994): the core (TIM23^CORE^) comprises three membrane-spanning proteins: Tim23, Tim17 and Tim50, which associates with different proteins to form complexes tailored for different tasks. Together with Tim21 and Mgr2 it forms the TIM23^SORT^ complex, capable of lateral release of proteins with hydrophobic sorting sequences. While association with the pre-sequence translocase-associated motor (PAM) forms the TIM23^MOTOR^ complex, responsible for matrix import.

Our current understanding of protein import *via* the TOM-TIM23^MOTOR^ complex is summarised in Figure 1A. After entry of the PCP through TOM, the electrical component of the proton-motive force (PMF) across the IMM – the membrane potential (Δψ; negative in the matrix) – is required, acting as an electrophoretic force on the positively charged pre-sequence (Martin et al., 1991)(Geissler et al., 2000)(Truscott et al., 2001). Δψ alone is sufficient for insertion of membrane proteins *via* the TIM23^SORT^ complex (Callegari et al., 2020), but complete import into the matrix by the TIM23^MOTOR^ complex requires an additional driving force: ATP hydrolysis by the main component of PAM, the mtHsp70 protein (Ssc1 in yeast)(Wachter et al., 1994), which pulls the rest of the PCP through to the matrix after the MTS has been imported. Finally, after crossing the IMM either way the MTS is cleaved by a matrix processing peptidase (MPP) (Vögtle et al., 2009).

**Figure 1:**
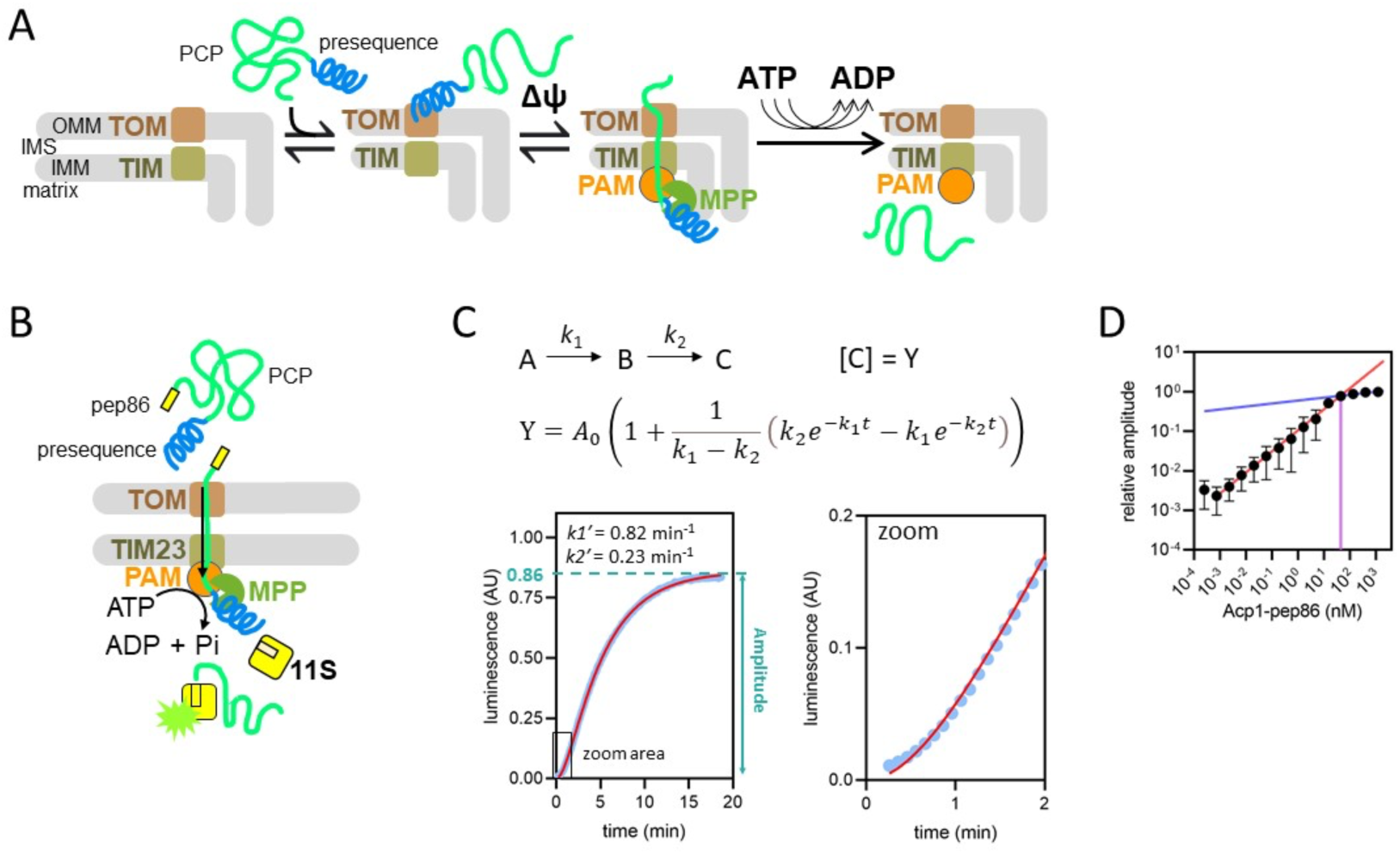
Model of PCP import into mitochondria and outline of the NanoLuc import assay. **A)** Simple model of presequence-containing precursor (PCP) import into mitochondria, showing binding of PCP to the TOM complex, Δψ-dependent movement of the presequence into the matrix and ATP-dependent translocation of the remainder of the protein. **B)** Diagramatic representation of the NanoLuc real-time import assay, which is essentially the model in A plus the binding of the C-terminal pep86 to internalised 11S which forms NanoLuc in the matrix. **C)** An example of luminescence data from the NanoLuc import assay of 1 µM DDL (length variant, see results) in energised mitochondria, showing the fit to a model for two consecutive, irreversible steps (see Methods). The final step gives rise to signal such that [C] (concentration of C) is proportional to luminescence. The order of the two steps is assigned arbitrarily. **D)** The effect of varying PCP concentration (Acp1-pep86) on amplitude of signal from import reactions. A straight line was fitted to the data where amplitude increased linearly with PCP concentration (red), and to the data where amplitude increased only marginally (blue). The intersect of these lines and corresponding PCP concentration (∼45 nM), the point of plateau, is also shown (purple).

The above model is primarily derived from end point measurements from a classical import assay involving autoradiography or Western blotting. However, this method is limited in its time resolution, and insufficient to provide a deep understanding of the individual steps that make up import, or their relative contributions to its kinetics. For this reason, we recently developed a highly time-resolved and sensitive assay which exploits a split NanoLuc enzyme (Pereira et al., 2019)(Dixon et al., 2015) to measure protein transport across membranes (Figure 1B). In the NanoLuc assay, PCPs tagged with a small fragment of the NanoLuc enzyme (an 11 amino acid peptide called pep86), are added to mitochondria isolated from yeast engineered to contain a matrix-localised large fragment of the enzyme (the enzyme lacking a single β-strand, called 11S). When the PCP-pep86 fusion protein reaches the matrix, pep86 binds rapidly and with tight affinity to 11S forming a complete NanoLuc luciferase. In the presence of the NanoLuc substrate (furimazine), this generates a luminescence signal proportional to the amount of NanoLuc formed. Luminescence is thus a direct readout of the amount of pep86 (and hence PCP) that has entered the matrix, up to the total amount of 11S. As expected, it is Δψ-dependent, affected by depletion of ATP, and sensitive to specific inhibitors of TIM23-dependent protein import (Pereira et al., 2019).

Here, we continue the use of the NanoLuc translocation assay to obtain precise, time-resolved measurements of protein delivery into the matrix mediated by the TOM and TIM23^MOTOR^ complexes. To add mechanistic detail to the above model (Figure 1A), we systematically varied the length and charge of the mature sequences of PCPs and profiled their import kinetics. To better understand the cause of any effects on the observable kinetic parameters (amplitude, rate and lag), we performed experiments under conditions where either of the two main driving forces, Δψ or ATP, had been depleted.

Our results suggest that IMM transport itself is fast in normally functioning mitochondria, and limited by the availability of Δψ. The rate of import is instead limited by transport across the OMM, which is strongly dependent on protein size, and initiation of transport across the IMM by the MTS. Analyses such as these, together with emerging structures of the import machinery eg. (Tucker and Park, 2019), will be fundamental to understanding the underlying molecular basis of mitochondrial protein import.

## RESULTS

### The import reaction is largely single turnover under the experimental conditions deployed

An exemplar NanoLuc import trace is shown in Figure 1C, collected using the model yeast matrix protein Acp1 (also used in previous import studies as a matrix-targeted precursor (Wurm and Jakobs, 2006)) fused to a pep86 (Acp1-pep86). The most intuitive parameter of this trace is amplitude (see below for full fitting details), which corresponds to the amount of NanoLuc formed when the reaction reaches completion, and thus the total number of import events; so long as the pep86 tag does not exceed matrix 11S. In order to verify that this was not the case we estimated the concentration of 11S in the mitochondria by quantitative Western blotting. An antibody raised against intact NanoLuc, capable of detecting 11S, was used to compare the quantities of mitochondrial 11S with known quantities of the purified protein (Figure 1 – figure supplement 1A). The results reveal high (µM) internal 11S concentrations with some variation between mitochondrial preparations (∼2.8 - 7.5 µM). We see no correlation between the amount of 11S and signal amplitude even with saturating PCP (Figure 1 – figure supplement 1B-C, and see below). Thus, we can conclude that the matrix concentration of 11S is in excess over the imported PCP, and not limiting the reaction, irrespective of how much is added to the outside.

We next measured signal amplitude over a wide range of concentrations of Acp1-pep86. Plotting the results shows that amplitude is linearly related to PCP concentration from 753 fM up to ∼ 45 nM, where it plateaus (Figure 1D). Because the mitochondrial matrix volume is only ∼1/12,000 of the total reaction volume (see Methods), if all 45 nM PCP were imported it would correspond to roughly 540 µM inside the matrix. This is not only far in excess of the internal 11S concentration (as low as ∼2.8 µM), but is also implausible simply from the amount of physical space available. Evidently then, only a tiny fraction of the PCP added reaches the matrix.

As neither the amount of PCP added, nor the amount of 11S in the matrix, appear to be limiting, we next tested to see whether the number of import sites might be having an effect. To estimate the number of import sites, we generated a PCP that can enter and give a signal, but which prevents subsequent import events through the same import site – *i*.*e*. forcing single turnover conditions. To do this, we fused dihydrofolate reductase (DHFR) to a model PCP; in the presence of the inhibitor methotrexate (MTX) DHFR folds tightly and cannot be imported (Pfanner et al., 1987)(Gold et al., 2017).

As expected, if DHFR is omitted (PCP-pep86) MTX has no effect (Figure 2A, grey bars), while if it is positioned N-terminal to pep86 (PCP-DHFR-pep86) we see very little luminescence with MTX present, consistent with blocked import (Figure 2A, purple bars). However, when DHFR is positioned C-terminal to pep86 (PCP-pep86-DHFR) with sufficient length between the two to span the TOM and TIM complexes (212 amino acids in this case, longer than the 135 required (Rassow et al., 1989)), we do see an import signal (Figure 2A, orange bars). This confirms that NanoLuc can form as soon as pep86 enters the matrix and does not require the entire PCP to be imported, as seen previously with the bacterial Sec system (Allen et al., 2020).

**Figure 2:**
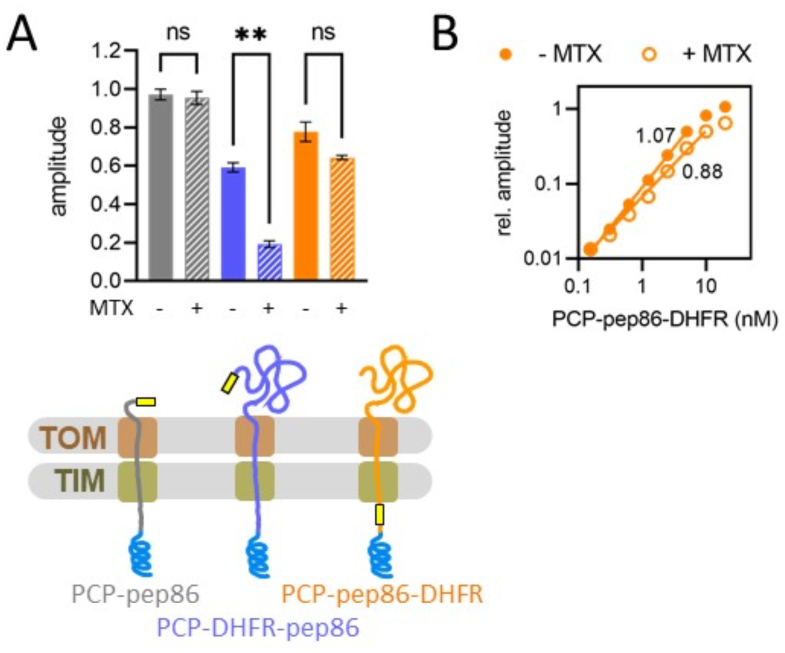
Basic characterisation of PCP import and turnover number. **A)** The effect of MTX on signal amplitude of three proteins (depicted schematically below): PCP-pep86 (grey), for which MTX should have no effect; PCP-DHFR-pep86 (blue), where MTX prevents entry of pep86; and PCP-pep86-DHFR (orange), where MTX limits import to one pep86 per import site. Bars show the average and SEM from three independent biological replicates. Differences between groups were analysed using a one-way ANOVA test, with Geisser-Greenhouse correction applied, followed by the Holm-Sidak multiple comparisons test. **, P<0.05; ns, not significant. **B)** Signal amplitude as a function of PCP-pep86-DHFR concentration in the absence (solid circles) and presence (open circles) of MTX.

Importantly, the presence or absence of MTX makes only a minor difference to the amplitude of this signal (Figure 2A). Indeed, the signal amplitude as a function of the [PCP-pep86-DHFR] is similar in the presence or absence of MTX (Figure 2B). The slope, which corresponds to the increase in amplitude per 1 nM PCP-pep86-DHFR, is 1.22 times greater in the absence of MTX, meaning only about 20% of the signal arises from turnovers beyond the first. Of course this does not mean that import is strictly single turnover – which would certainly seem implausible for fully functional mitochondria in their native environment – it does suggest that it behaves as single turnover under the conditions here using isolated mitochondria (without the cytosol).

It has previously been shown that signal amplitude can be reduced by depleting Δψ (Pereira et al., 2019), which would suggest that available energy limits protein import. This can be reconciled with the apparent single turnover nature of the reaction if ‘resetting’ the channel after import – possibly through dimerisation of TIM23, as previously reported (Bauer et al., 1996) – requires additional energy input.

### Kinetic analysis of import suggests two major rate-limiting steps

In addition to the amplitude data, the import traces contain information about the kinetics of the reaction. Looking again at the data in Figure 1C, it can be seen that import does not start at its maximum rate; rather there is a lag before import accelerates. This is characteristic of reactions with multiple consecutive steps, where only the last one gives rise to a signal. As an approximation, the data fitted well to an equation for a two-step process where the second gives rise to the signal (A→B→C, see also Methods), which gives two apparent rate constants (*k*_1_′ and *k*_2_′) in addition to amplitude (Fersht, 1984). Close inspection of the fit (Figure 1C, right panel) suggests that adding additional steps would marginally improve the fit, however these additional rate constants would be fast and poorly defined; two steps therefore represents a reasonable compromise between accuracy and complexity.

In the simplest case possible, where the two steps are irreversible and have very different values, *k*_1_′ and *k*_2_′ correspond to the two rates for these steps (*k*_1_ and *k*_2_)(Fersht, 1984). This is complicated if the reactions are reversible (in which case the reverse rates also factor), or if *k*_1_ and *k*_2_ are very similar (where they are both convoluted into *k*_1_′ and *k*_2_′). Nonetheless, this analysis is very useful for understanding the mechanism of import (see below) – especially under conditions where *k*_1_ and *k*_2_ are well separated.

It should be noted that, because we have no information for the concentration of the intermediates, the order of the two steps cannot be determined *a priori*. However, as detailed below, they can be distinguished by perturbing the system and seeing how this affects the different rates. From this, and based on the results in the following sections, we assign *k*_1_′ as transport of the PCP through TOM and *k*_2_′ as subsequent engagement of the MTS with TIM23. It is also important to note that any additional step faster than about 5 min^-1^ will not be resolved in our experimental set up using a multi-plate reader (see detailed explanation in Figure 1 – figure supplement 2A), and will instead manifest as a small apparent lag before the signal appears (equal to 1/*k*_step_, where *k*_step_ is the rate constant for that process)(Allen et al., 2020). This includes formation of NanoLuc: it is >7.4 min^-^ _1_ even at the lowest estimated 11S concentration, as determined in solution (Figure 1 – figure supplement 2B), and the import kinetics are not appreciably affected by the internal concentration of 11S as noted above; Figure 1 – figure supplement 2C.

### Import is dependent on total protein size

To begin to validate what the two apparent rates correspond to, we first designed and purified two series of four PCPs, varying either in total length or in the N- to C-terminal positioning of pep86 (Figure 3A). The length variants all similarly contained the pre-sequence of Acp1 followed by the Acp1 mature domain, with pep86 (L) at the C-terminus. Increase in length was achieved by repeating the mature part of Acp1 up to three times. In between each Acp1 mature domain we included a scrambled pep86 sequence (D), which does not interact with 11S (Allen et al., 2020), such that each tandem repeat has the same overall amino acid (aa) composition.

**Figure 3:**
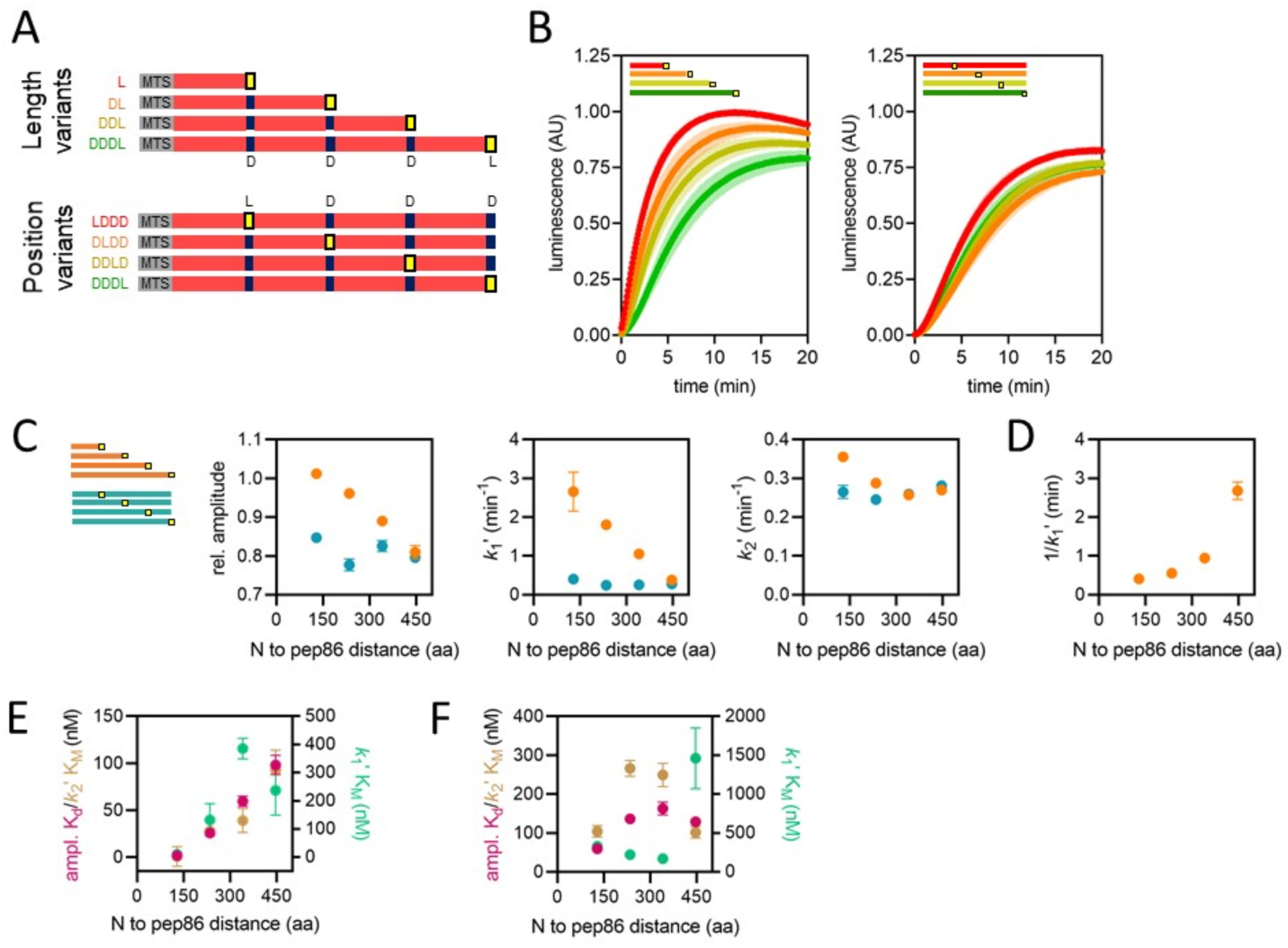
Using proteins of varying lengths to elucidate import kinetics. **A)** Schematic of two protein series (length variants and position variants), with native MTS and mature part of Acp1 in grey and red respectively, pep86 in yellow (L for Live) and scrambled pep86 in dark blue (D for Dead, i.e. it does not complement 11S). **B)** Example of import traces for length variants (left panel) and position variants (right panel). Error bars shown partially transparent in the same colours as the main traces. Those smaller than the main trace are not shown. SD from biological triplicate, each conducted in duplicate. **C)** Parameters obtained from two step fits to the data shown in panel B. The length variant series is shown in orange and the position variant series in teal. Error bars show SEM from three independent biological experiments, each conducted in duplicate. Error bars smaller than symbols are not shown. **D)** Reciprocal of *k*_1_′ as a function of PCP length (same data as in panel C) – i.e. the time constant for that steps – for the length variants. **E)** The concentration dependence of length variants. Secondary data from import assays with varying concentrations of length series proteins (4-6 independent biological replicates) were fitted to the Michaelis-Menten equation, from which apparent K_d_s and K_M_s are derived. Error represents the SEM of this fitting. **F)** As in panel F but with the position variant proteins.

The length variant set was designed to reveal PCP size-dependence of any import step. The other set (position variants) were all identical to the longest length-variant PCP (four tandem repeats), but with the active pep86 (L) in different positions. Because the position variants (abbreviated as LDDD, DLDD, DDLD and DDDL) are identical save for the number of amino acids that must enter the matrix before the NanoLuc signal arises, all transport steps (including passage through TOM) should be the same for the whole set. Any differences in their import kinetics must therefore arise from the time it takes them to pass through TIM23, and not the steps prior to that. Note that as shown above (Figure 2A) and previously (Allen et al., 2020), localisation to an internal loop does not compromise the ability of pep86 to interact with 11S.

Import of all four length variants (L, DL, DDL and DDDL) and position variants (LDDD, DLDD, DDLD and DDDL) at high concentration (1 µM, which is saturating for all parameters, see below (Figure 3 – figure supplement 1)) fit well to the simple two-step model, giving an amplitude and two apparent rate constants, with the faster one assigned as *k*_1_′ and the other as *k*_2_′. Import traces and the results of fits to the two-step model are plotted in Figure 3B and C respectively. We observe no significant difference between any of the four position variants with respect to any of the three parameters, indicating that transport through TIM23 is fast, and does not contribute appreciably to the kinetics of import.

For the length series, signal amplitude is inversely correlated with protein length (Figure 3C, left panel in orange). Let us suppose that, at any point during processive translocation, an import site can become compromised; for instance, by a PCP becoming trapped in the channel. In this scenario it would be reasonable to expect a longer protein to have a higher chance of failing to reach the matrix. But if this were the cause of the dependence of signal amplitude on protein length, we would expect a similar dependence for the position variants, which is not the case (Figure 3C, in teal). Instead it seems that small proteins are able to accumulate at higher levels in the matrix compared to large ones. This is consistent with our previous conclusion that the amount of 11S does not limit import, as this would result in the same maximum amplitude for all proteins.

Strikingly, we find that *k*_1_′ has a strong inverse correlation with PCP length (but not pep86 position), *i*.*e*. it is faster for smaller proteins (Figure 3C, middle panel). The most likely explanation for this is that *k*_1_′ corresponds to transport of the entire length of the protein across a membrane. Even more surprisingly, the corresponding step time (1/*k*_1_′) increases not linearly but exponentially as a function of PCP length (Figure 3D). This means that longer PCPs complete step *k*_1_′ more slowly *per amino acid*. Exponential length-dependence is not a characteristic of a powered or biased directional transport, such as we have seen previously for the Sec system (Allen et al., 2020), but rather an unbiased reversible diffusion-based mechanism (Simon et al., 1992). For *k*_2_′, meanwhile, there is little difference between the variants; indeed, with the exception of L, good fits can be obtained when *k*_2_′ is fixed globally (Figure 3C, right panel). Unlike *k*_*1*_′ therefore, *k*_2_′ probably corresponds to something other than transport across a membrane.

#### Concentration dependence of the two major rate-limiting steps of import

A simple way to assign rate constants to specific events is to measure their dependence on concentration: only steps that involve association between PCP upon the initial contact with the import machinery (with the TOM complex) should show any concentration effect. We therefore measured protein import for both the length and position variants over a range of PCP concentrations ([PCP]), and fitted the data for each concentration to the two-step model. Next, we plotted the concentration dependence of each of the three resulting parameters (Figure 3 – figure supplement 1), and fitted them to a weak binding (amplitude) or Michaelis Menten (k1′ and k2′) equation (Figure 3E-F). It should be noted that the KM values are rough estimates only, as k1′ and k2′ are difficult to assign precisely.

Unexpectedly, all three parameters show a dependence on [PCP] for the length series. The apparent K_M_s for *k*_1_′, (Figure 3E, teal) are in the low 100s of nM and not systematically dependent on [PCP] – both reasonable for initial association of PCP and TOM. The K_d_s for amplitude and K_M_s for *k*_2_′, meanwhile (magenta and brown, respectively in Figure 3E), are very similar to each other: they are very low (tight affinity), but increase with increasing PCP length. Because amplitude and *k*_2_′ behave identically, it seems reasonable to assume that they reflect the same process, i.e. the final kinetic step of transport (because amplitude is by definition successful transport). The precursor length-dependence means that, effectively, longer PCPs require a higher concentration to reach maximum amplitude (Figure 3E), even though that amplitude is lower (Figure 3B-C). One possible explanation for this is that at very low PCP concentrations, affinity of PCP in the IMS for TIM23 begins to become limiting. Just as before, we find no systematic difference between the position variants (Figure 3F) – again suggesting that passage of the PCP through TIM23 is not limiting the overall import rate.

### Depletion of Δψ and ATP have very different effects on import

The two driving forces (Δψ and ATP) act at different stages of import (Figure 1A), so to help assign *k*_1_′ and *k*_2_′ we depleted each and measured import of the length and position variants. Partial depletion of Δψ by pre-treatment of mitochondria with 10 nM valinomycin, a potassium ionophore, causes a decrease in signal amplitude for all length and position variants, affecting them roughly equally (Figure 4A, left panel). Valinomycin treatment also affects both the apparent rate constants: *k*_1_′ is somewhat slowed for shorter proteins but largely unaffected for longer ones (Figure 4A, middle panel), while *k*_2_′ is somewhat slowed for short proteins but dramatically reduced for longer ones (Figure 4A, right panel).

**Figure 4:**
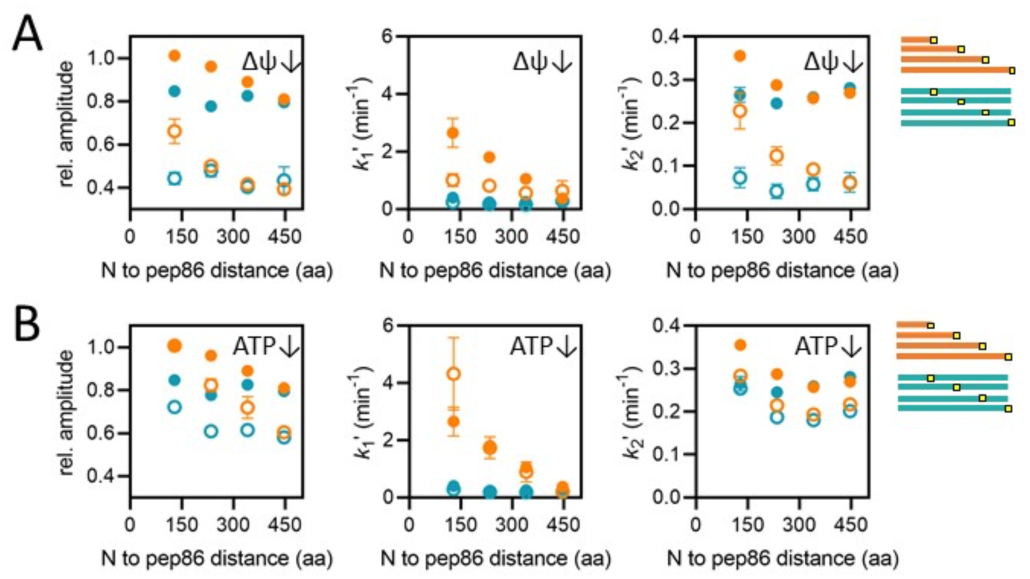
Effects of energy depletion on import of the length and position variants. **A)** Import in the presence (solid circles) or absence (open circles) of Δψ, for the length (orange) and position (teal) series. Depletion of Δψ was achieved by a 5 min pre-treatment of mitochondria with 10 nM valinomycin. Plots show amplitude (left), *k*_1_′ (middle) and *k*_2_′ (right) extracted from two-step fits to import traces as a function of PCP length or pep86 position. Each point is the average and SEM of three independent biological replicates. **B)** As in panel A, but without (solid circles) or with (open circles) ATP depletion instead of valinomycin. Matrix ATP was depleted by excluding ATP and its regenerating system from the assay mix (see results section for full description).

Depletion of matrix ATP was achieved simply by excluding ATP and its regenerating system from the assay buffer. Endogenous matrix ATP under these conditions is minimal, as is evident from the fact that import becomes highly sensitive to antimycin A, an inhibitor of oxidative phosphorylation (Figure 4 – figure supplement 1). This sensitivity arises because ATP is required for hydrolysis by the ATP synthase to maintain Δψ in the absence of oxidative phosphorylation (Campanella et al., 2008). Import experiments performed with depleted ATP show reduced amplitude, but unlike valinomycin this effect is more pronounced for the longer PCPs (Figure 4B, left panel) – consistent with proposed role for ATP in promoting transport of the mature part of the PCP. ATP depletion has little or no effect on *k*_1_′ (Figure 4B, middle panel), and a relatively minor effect on *k*_2_′ (Figure 4B, right panel), affecting both the length and position variants roughly equally.

### A simple working model for import based on the above results

Taking all the above observations together, we can as alluded to earlier propose a simplified model for import that incorporates two major rate-limiting steps. Based on its dependence on PCP concentration (Figure 3E) we assign *k*_1_′ as dependent on the initial interaction between PCP and TOM. However this concentration dependence saturates with an apparent K_M_ of around 100-200 nM. Such saturating behaviour suggests a rapid binding equilibrium followed by a slower step (just as in Michaelis Menten kinetics), i.e.:

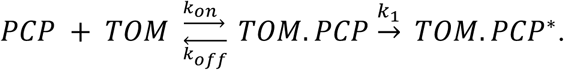

The strong dependence of *k*_1_′ on PCP length (Figure 3C, middle panel) provides a clue as to the nature of *k*_1_ – it is likely to correspond to passage of the PCP across the OM, through the TOM complex. The non-linear dependence of step time (1/*k*_1_′) on PCP length (Figure 3D) also suggests that this step is at least partially diffusional, rather than driven by an active energy-dependent directional motor. Furthermore, it suggests that, under these experimental conditions at least, the entire PCP passes through TOM before transport through TIM23 is initiated.

The second rate constant, *k*_2_′ is somewhat sensitive to ATP (Figure 4B, right panel), and so most likely comes at the end of import, as the contribution of Hsp70 requires at least some of the PCP to be in the matrix. Since *k*_2_′ shows very little dependence on PCP length in energised mitochondria (Figure 3C, right panel), we propose that it is primarily the Δψ-dependent insertion of the pre-sequence through TIM23, not the subsequent passage of the unfolded passenger domain that is limiting (although both presumably contribute to the apparent rate constant). However, under conditions of Δψ depletion, a length-dependence of *k*_2_′ emerges (Figure 4A, right panel): this is consistent with import rate of the rest of the PCP being affected by Δψ ((Schendzielorz et al., 2017), and see also below). It is also possible that transport of longer PCPs has a higher chance of failure, with the PCP slipping back into the IMS – this would be a useful mechanism to prevent TIM23 complexes becoming blocked with mis-folded/compacted PCPs, and would explain the difference in the effect of Δψ depletion on the length and position variants.

Putting all of this together, we propose the following minimal kinetic scheme for PCP import:

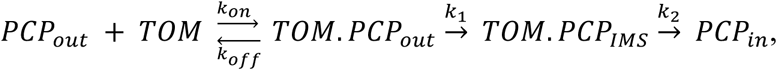

where the subscript to PCP indicates its location (<u>out</u>side the OM, in the <u>IMS</u>, or <u>in</u>side the matrix). In this model, *k*_on_ and *k*_off_ are both fast compared with *k*_1_, and give an affinity (K_d_ = *k*_off_/ *k*_on_) in the order of 100 nM, similar to the affinity of a bacterial secretion preproteins to bacterial inner membrane vesicles (Hartl et al., 1990). The two extracted rate constants can be approximately determined as ([PCP] designates PCP concentration):

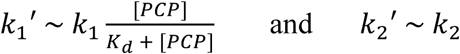

This model fits the data, and we believe it is the most reasonable interpretation of the above experimental results. However it still leaves open several questions, notably the extent to which *k*_1_ and *k*_2_ are reversible. For example, the fact that *k*_1_′ is somewhat affected by valinomycin (Figure 4A, middle panel) suggests that *k*_1_ is reversible. Given that passage through TOM can occur in the absence of Δψ (Mayer et al., 1993)(Lill et al., 1992), slowing *k*_2_ would then leave more opportunity for diffusion back out of the IMS through TOM, a process that occurs in the absence of ATP (Ungermann et al., 1996). In addition, we cannot determine from this data exactly at what stage handover from TOM to TIM23 occurs. The results suggest that PCP passes through TOM completely before engaging with TIM23, but it is not clear whether this is a necessary part of the mechanism or merely an effect of the relative rates under these conditions. Nor can we determine whether handover from TOM to TIM23 is direct, or if the PCP can dissociate from TOM before binding to TIM23.

### Changing PCP net charge affects import amplitude and rate differently

Δψ, the electrical component of the PMF (positive outside), has been proposed to act primarily upon positively charged residues in the PCP, pulling them through electrophoretically (Martin et al., 1991)(Geissler et al., 2000)(Truscott et al., 2001). To test this idea, we designed a series of proteins, based on a engineered version of a classical import substrate: the N-terminal section of yeast cytochrome *b*_2_ lacking the stop-transfer signal (Δ43-65) to enable complete matrix import (Gold et al., 2014). The variant PCPs differed only in the numbers of charged residues (Figure 5A); of the same length (203 amino acids), but spanning 5.43 units of pI ranging from 4.97 to 10.4. Import of these charge variants under saturating conditions (1 µM PCP) was measured using the NanoLuc assay as above and representative traces are shown in Figure 5B (with complete data in Figure 5 – figure supplement 1).

**Figure 5:**
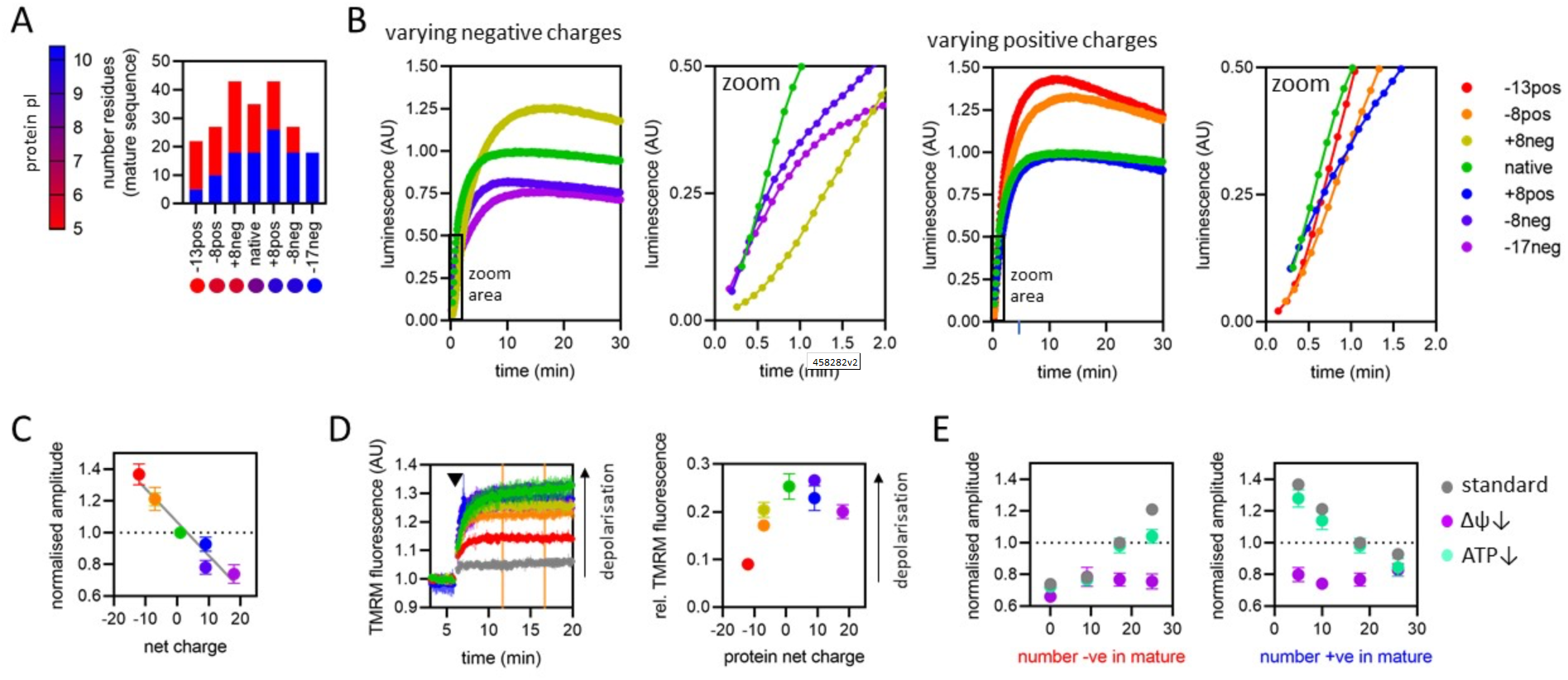
The effect of PCP charge on import kinetics. **A)** Overview of the charge variant protein series, showing numbers of positively (blue) and negatively (red) charged residues in the mature part of each protein, and symbols for each protein with colours corresponding to theoretical pI, according to the scale shown on the left. All proteins in the charge variant series have the same length (203 amino acids), and are based on the N-terminal section of yeast cytochrome *b*_2_ lacking the stop-transfer signal (Δ43-65) to enable complete matrix import (Gold et al., 2014). **B)** Import traces for the charge variant proteins in which the number of negative (left) and positive (right) charges are varied, normalised to the native PCP, coloured by rainbow from most negative (red) to most positive (violet). Data shown are a single representative trace; this is because starting points for each data set are slightly offset due to the injection time of the plate reader. Full data – three biological replicates each performed in duplicate – are shown in Figure 5 – figure supplement 1. **C)** Amplitudes obtained from panel (**B**) as a function of net charge (coloured as in panel B), with a line of best fit shown. The data point for the +8neg protein (yellow) is in the same position as the -8pos protein (orange) and is mostly hidden. **D)** TMRM fluorescence over time in isolated yeast mitochondria (left), with PCPs added at the time indicated by arrowhead. A no protein control (buffer only) is shown in grey, and the remaining traces are shown with the PCP coloured as in panel B. Average TMRM fluorescence over a 5 minute window (between orange vertical lines) was calculated for each trace then plotted, relative to no protein control, against protein net charge (right). Data shown is mean ± SD from three biological repeats. **E)** Amplitude (normalised to the native PCP in standard conditions) of import signal for the charge variants, where numbers of negatively (left) or positively (right) charged residues is varied, under standard reaction conditions (grey) or when Δψ (purple) or ATP (green) is depleted. Each data point is the mean ± SEM from three biological repeats (shown in Figure 5 – figure supplement 1B-C). Error bars smaller than symbols are not shown.

The most immediately striking observation is that amplitude is strongly inversely correlated with net charge of the PCP – *i*.*e*. the opposite of what might be expected given the direction of Δψ (Figure 5C). To understand why this would be, we turned to our earlier interpretation of signal amplitude: that it is limited by the availability of Δψ. If transport of positively charged residues depletes Δψ while transport of negatively charged residues replenishes or maintains it, this could explain why negatively charged proteins accumulate to a higher level.

To test this hypothesis, we monitored Δψ in isolated mitochondria over time by measuring TMRM fluorescence, then assessed the effect of adding the PCPs with differing net charge (Figure 5D). The PCPs did indeed cause strong depletion of Δψ and, moreover, this effect diminished with increasing net negative charge. Increasing net positive charge did not seem to result in enhanced depletion of Δψ, but TMRM does not resolve Δψ well in this range, so this does not necessarily mean that this effect is not occurring. A second prediction from this hypothesis is that membrane depolarisation prior to protein import will abolish the correlation between net charge and amplitude. This is indeed exactly what we observe: valinomycin reduces amplitudes for all PCPs, but the effect is greater for more negatively charged PCPs, bringing all amplitudes to about the same level (Figure 5E). Depleting ATP, meanwhile, has very little effect on amplitude, just as for the Acp1-based PCPs.

It is also clear, from the import traces for the charge series, that positively charged PCPs are imported much faster than negatively charged ones (albeit reaching a lower final amplitude; Figure 5B-C). This is again consistent with Δψ specifically assisting the transport of positively charged residues (Martin et al., 1991)(Geissler et al., 2000)(Truscott et al., 2001). Unlike the length variants based on Acp1 (Figure 3A), however, not all of the import traces from the charge variants (Figure 5B) fit to the two step model (see Methods and Figure 1C). While the more negatively charged ones have a clear lag before reaching their maximum rate, the positively charged ones appear to have only a single rate-limiting step, or even to have a burst of rapid import, followed by a slower phase (Figure 5B). Because steps are only resolved on the plate reader if they are ≤ about 5 min^-1^, the most likely explanation for this is that one step has become too fast to measure. This is most likely transport through TIM23, which is strongly Δψ-dependent and thus presumably faster for more positively charged proteins. A burst suggests multiple turnovers, not seen for the Acp1-based DHFR-pep86 constructs (Figure 2), with the first one very fast and subsequent ones limited by a slower resetting of TIM23 (see Discussion).

### Validation of the observed charge and size effects with native PCPs

While the use of artificial PCPs, as above, allows their properties to be varied in a systematic manner, it is possible that these modifications will affect native features with fundamental roles in the import process. To confirm that the above observations hold true for native PCPs we performed import experiments with four pep86-tagged native PCPs differing in length and charge. We chose the F_1_ α and F_1_ β subunits of the mitochondrial ATP synthase, both large proteins (>500 amino acids) with mature amino acid sequences differing in predicted pI by ∼ 1.55 (F_1_ β = 5.43 and F_1_ α = 6.98); and two smaller proteins (<200 amino acids), Acp1 and Mrp21, with predicted mature sequence pIs of 4.87 and 10.00 respectively (Figure 6A).

**Figure 6:**
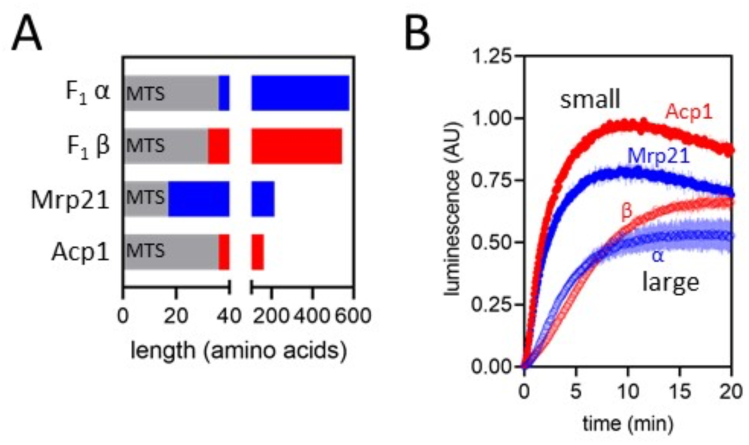
Import of pep86 fused native precursors. **A)** Schematic representation of the four native PCPs chosen: F_1_α (long, positively charged, predicted pI of mature part is 6.98), F_1_β (long, negatively charged, predicted pI of mature part is 5.43), Mrp21 (short, positively charged, predicted pI of mature part is 10.00) and Acp1 (short, negatively charged, predicted pI of mature part is 4.87). **B)** Import traces for the four PCPs in panel A under standard conditions (1 µM PCP), normalised to Acp1. Each trace is the mean ± SD of three biological repeats.

Consistent with our earlier results, we see higher amplitudes for the shorter and more negatively charged PCPs (Figure 6B), and faster import of the shorter PCPs than the longer ones (Figure 6B). The effect of net charge holds true for the larger PCPs, which both have clear two-step import (Figure 6B), but the small PCPs appear to have only a single rate-limiting step, and do not differ significantly in import rate (Figure 6B). Presumably the charge dependence only becomes measurable when transport through TIM23 is slow enough to be appreciable. Overall, these results suggest that the data collected with artificial PCPs will hold true for native ones as well.

## DISCUSSION

Protein import into mitochondria is, by nature, a complicated process with machineries in two membranes having to coordinate with one another as well as with parallel import pathways to deliver a wide range of proteins to their correct destinations. Here, we have built a minimal mechanistic model of one of the major import routes – the TOM-TIM23^MOTOR^ pathway of matrix proteins – using a high-resolution import assay based on NanoLuc (Pereira et al., 2019). Our results suggest that two major distinct events are responsible for the majority of the PCP transit time: passage of the PCP through the TOM complex and insertion of the pre-sequence through the TIM23^MOTOR^ complex. By contrast, the initial binding of PCP to TOM is fairly rapid, as is passage of the mature PCP domain through TIM23. Crucially, the rates of the different steps correlate very poorly with the amount of PCP in the matrix when the reaction ends, which has always been the conventional readout of import. It therefore seems that this pre-steady-state kinetic approach will be critical in the future, both for further dissecting import via the TOM and TIM23^MOTOR^ complexes and for understanding the other pathways that together comprise the mitochondrial protein import machinery.

Import appears to be largely single turnover under our experimental conditions, that is each import site only imports a single PCP. While this is fortuitous in that it allows us to access pre steady-state events easily, it is incongruent with mitochondrial protein import *in vivo*. Nonetheless, this almost certainly holds true for decades of experiments using the classic method, and offers an explanation as to why these methods require such high concentrations of mitochondria for detection of import. We propose that, under experimental conditions, import is limited by the amount of energy available in the form of Δψ. Indeed, measurements of Δψ using TMRM confirm that PCP import causes a depolarisation of the IMM that is not restored. Also consistent with Δψ being consumed, we find that the PCPs that require more total energy to import (such as longer ones), or that are likely to consume more Δψ (positively charged ones) reach a lower concentration in the mitochondrial matrix. Presumably isolated mitochondria, while capable of respiration and ATP synthesis, do not have the full restorative powers available to those inside cells.

The mechanism by which Δψ-depletion leads to single turnover conditions may relate to the requirement of Δψ for dimerization of TIM23 and recruitment of Tim44, both required for delivery to the matrix (Bauer et al., 1996)(Martinez-Caballero et al., 2007)(Demishtein-Zohary et al., 2017)(Ting et al., 2017)(Ramesh et al., 2016). As PCPs bind only to TIM23 complexes containing two Tim23 subunits and, during transport, disrupt this conformation, loss of Δψ would prohibit the resetting of the TIM23 complex to allow further turnovers after the first one (Bauer et al., 1996). With some of the faster importing PCPs we do indeed see a rapid burst of import followed by a slower phase, as would be expected for multiple turnovers where the first is fast. This could therefore provide an opportunity for future studies to investigate this priming event.

Previous studies have shown that the TOM complex is in excess over TIM23, with 1 mg yeast mitochondria containing ∼17-20 pmol TIM23 (∼9-10 pmol dimer) and estimations of between 85 and 250 pmol TOM40 (Sirrenberg et al., 1997)(Dekker et al., 1997). In our experiments, this TIM23 dimer concentration equates to ∼62.5 fmol per well (10 pmol.mg^-1^ × 50 µg.ml^-1^ × 125 µl) – similar to the estimated amount of 11S (∼28-76 fmol per well, based on an estimated 4.46-12.17 pmol.mg^-1^). This close correspondence presumably explains why we find that 11S is not limiting, but intriguingly, it also suggests that each import site only imported on average one 11S, even though 11S import occurred in live yeast before mitochondrial isolation. This correspondence may not be coincidental; if the number of TIM23 sites limited import, this could be calibrated as a regulatory mechanism to avoid matrix-derived proteotoxic stress.

The transfer of PCPs from TOM to TIM23 is thought to involve cooperative interactions of subunits of the two complexes (Gomkale et al., 2021)(Callegari et al., 2020). But the extent to which transport of PCPs across the OMM and IMM is coupled *in vivo*, remains unknown. It has been suggested that the rate of PCP passage through the OMM is one factor that determines whether PCPs are transferred to the matrix or released laterally into the IMM (Harner et al., 2011b), implying simultaneous and cooperative activities of TOM and TIM23. PCPs have been captured spanning both membrane complexes at the same time in super-complexes of ∼600 kDa (Gomkale et al., 2021)(Dekker et al., 1997)(Gold et al., 2014)(Chacinska et al., 2010), suggesting that import through TOM does not have to be complete before import through TIM23 can begin.

Contrasting with this, however there is also evidence to suggest that the TOM and TIM23 complexes can transport PCPs independently, in steps that are not necessarily concurrent. Matrix import of PCPs has been observed in mitoplasts (Hwang et al., 1989)(Ohba and Schatz, 1987), in which the OMM has been removed, suggesting that a handover from TOM is not absolutely required. Furthermore, the *in vivo* existence of TOM-TIM23 super-complexes is unconfirmed. They have been detected only when engineered PCPs with C-terminal domains that cannot pass through TOM are used (Chacinska et al., 2003), and only under these artificial conditions do TOM and TIM23 subunits co-immuno-precipitate or co-migrate on native polyacrylamide gels (Horst et al., 1995). Perhaps their assembly is more dynamic and transient, relying on other OMM-IMM contact sites such as the MICOS complex (von der Malsburg et al., 2011)(Hoppins et al., 2011)(Harner et al., 2011a). Moreover, the N-terminal domain of Tim23, which tethers the IMM and OMM, is not required for either PCP import though TIM23, or TOM-TIM23 super-complex formation (Chacinska et al., 2003).

Our results also hint that this handover is not absolutely required. The data here suggest that transport of a PCP through TOM is reversible, and therefore possible in the absence of TIM23 activity. Reverse transport of proteins through TOM, and in some cases also through TIM23, has been observed previously, although this process is not well understood. For example, proteins that are reduced or conformationally unstable in the IMS can retro-translocate to the cytosol *via* TOM40, and the efficiency of this process is relative to protein size (both linear length and 3D complexity); smaller proteins are more efficiently retro-translocated (Bragoszewski et al., 2015). Notably, under physiological conditions, PINK1 is cleaved in the IMM by PARL, releasing the C-terminal region for release back to the cytosol for proteosomal degradation. But the process is not well understood, such as if, and how, it is regulated, and if a driving force is required. Additionally, we see some PCP concentration dependence of *k*_2_′; if direct interaction of TOM with TIM23 were strictly required then *k*_2_ would not be affected by PCP concentration, but if PCP can accumulate in the IMS this would explain our finding.

Overall, the above analysis provides good estimates of the two rate-limiting steps for import, and provides evidence as to the constraints that act upon the other (non-rate-limiting) steps. If a few of the above questions are resolved, we believe it should be possibly to construct a complete kinetic model of mitochondrial import, as has been recently achieved for the bacterial Sec system (Allen et al., 2020).

## MATERIALS AND METHODS

### Strains and plasmids

*E. coli* α-select cells were used for amplifying plasmid DNA and BL21 (DE3) used for protein expression. Genes encoding pep86 (trademarked as ‘SmBiT’ (Dixon et al., 2015)) -tagged mitochondrial PCP proteins (from MGW Eurofins or Thermo Fisher Scientific) were cloned into either pBAD, pRSFDuet or pE-SUMOpro. YPH499 yeast cell clones transformed with pYES2 containing the mt-11S gene under control of the GAL promoter, used previously (Pereira et al., 2019), were used for isolation of mitochondria containing matrix-localised 11S (trademarked as ‘LgBiT’ (Dixon et al., 2015)). *E. coli* cells were routinely grown at 37°C on LB agar and in either LB or 2xYT medium containing appropriate antibiotics for selection. Yeast cells were grown at 30°C on synthetic complete dropout (Formedium) agar supplemented with 2% glucose, penicillin and streptomycin, or in synthetic complete dropout medium, supplemented with 3% glycerol, penicillin and streptomycin in baffled flasks. For yeast cells with mitochondrial matrix-localised 11S, mt-11S was expressed by adding 1% galactose at mid-log phase, 16 hours prior to harvesting of cells.

### Protein production and purification

BL21 (DE3) cells from a single colony, containing the chosen protein expression plasmid were grown in LB overnight then sub-cultured in 2XYT medium until OD_600_ reached 0.6. For pBAD and pRSFDuet plasmids protein expression was induced by adding arabinose or IPTG respectively. Cells were harvested 2-3 hours later and lysed using a cell disrupter (Constant Systems Ltd.). Proteins were purified from inclusion bodies using Nickel affinity chromatography on prepacked HisTrap FF columns (Cytiva, UK), followed by ion exchange chromatography on either HiTrap Q HP or HiTrap SP HP columns (Cytiva, UK) depending on protein charge, described in full previously (Pereira et al., 2019). Proteins from pE-SUMOpro plasmids (those containing DHFR domains), were expressed by adding IPTG, and cells harvested after 18 hours of further growth at 18°C. Proteins were purified at 4°C from the soluble fraction, essentially as before (Aelst et al., 2019), but with 250 mM NaCl in their “Buffer C”. A further purification step, on a HiLoad 16/60 Superdex gel filtration column (Cytiva, UK) was included to remove remaining contaminants. A full list of PCPs, their amino acid sequences and respective expression vectors are given in Supplementary Table 1.

### Isolation of mitochondria from yeast cells

Yeast cells were harvested by centrifugation (4,000 x g, 10 min, room temperature) and mitochondria isolated by differential centrifugation (Daum et al., 1982). Briefly, cell walls were digested with zymolyase in phosphate-buffered sorbitol (1.2 M sorbitol, 20 mM potassium phosphate pH 7.4), after being reduced with DTT (1 mM DTT in 100 mM Tris-SO4 at pH 9.4, for 15 min at 30°C). Cells were disrupted at 4°C with a glass Potter-Elvehjem homogeniser with motorised pestle in a standard homogenisation buffer (0.6 M sorbitol, 0.5% (w/v) BSA, 1 mM PMSF, 10 mM Tris-HCl pH 7.4). The suspension was centrifuged at low speed (1,480 x *g*, 5 min) to pellet unbroken cells, cell debris and nuclei, and mitochondria harvested from the supernatant by centrifugation at 17,370 x *g*. The pellet, containing mitochondria, was washed in SM buffer (250 mM sucrose and 10 mM MOPS, pH 7.2), and then centrifuged at low speed again, to remove remaining contaminants. The final mitochondrial sample, isolated from the supernatant by centrifugation (17,370 x *g*, 15 min), was resuspended in SM buffer and protein quantified by bicinchoninic acid (BCA) assay (Smith et al., 1985) using a bovine serum albumin protein standard. Mitochondria were stored at -80°C, at a concentration of at 30 mg/ml in single use aliquots, after being snap frozen in liquid nitrogen.

### Western blotting

Samples of mitochondria from yeast cells were solubilised in SDS-PAGE sample buffer (2 % (w/v) SDS, 10% (v/v) glycerol, 62.5 mM Tris-HCl pH 6.8, 0.01% (w/v) bromophenol blue, and 25 mM DTT), and fractionated on a 15% (w/v) acrylamide, 375 mM Tris pH 8.8, 0.1% (w/v) SDS gel with a 5% (w/v) acrylamide, 126 mM Tris pH 6.8, 0.1% (w/v) SDS stacking gel, in Tris-Glycine running buffer pH 8.3 (25 mM Tris. 192 mM glycine, 0.1% (w/v) SDS). Proteins were electro-transferred to PVDF membrane in 10 nM NaHCO_3_, 3mM Na_2_CO_3_, then membranes incubated in blocking buffer (TBS (50 mM Tris-Cl pH 7.5, 150 mM NaCl) containing 0.1% (v/v) Tween 20 and 5% (w/v) skimmed milk powder). 11S protein was detected with a rabbit polyclonal antibody from Promega, and Tom40 with a rabbit polyclonal antibody produced by Cambridge Research Biochemicals (Billingham, UK). Primary antibody incubations were at 4°C for 18 h in blocking buffer. Membranes were washed in TBS containing 0.1% (v/v) Tween 20, three times, each for 10 minutes, before incubation for 1 hour with a HRP-conjugated goat secondary antibody against rabbit IgG (Thermo Fisher Scientific), in blocking buffer. Membranes were washed, as before, and antibodies visualised using 1.25 mM luminol, with 198 µM coumaric acid as enhancer, and 0.015% (v/v) H_2_O_2_ in 100 mM Tris-Cl pH 8.5.

### NanoLuc import assay

Unless stated otherwise, import experiments were performed at 25°C with mt-11S mitochondria diluted to 50 µg/ml in import buffer (250 mM sucrose, 80 mM KCl, 1 mM K_2_HPO_4_/KH_2_PO_4_, 5 mM MgCl_2_, 10 mM MOPS-KOH and 0.1% (v/v) Prionex reagent (Merck), pH 7.2), supplemented with 2 mM NADH, 1 mM ATP, 0.1 mg/ml creatine kinase, 5 mM phosphocreatine, and 1 µM pep86-tagged PCP protein. We also added 10 µM GST-Dark protein; a fusion of glutathione S-transferase and a peptide with high affinity for 11S that inhibits pep86 binding and concomitant enzymatic activity, and thereby reduces background signal caused by trace amounts of 11S outside the mitochondrial matrix (Pereira et al., 2019). Mitochondria and GST-Dark were added to 1X import buffer at 1.25X final concentrations (mixture 1), and pep86-tagged PCP, NADH, ATP, creatine kinase and phosphocreatine added to 1X import buffer at 5X final concentrations (mixture 2) so that import reactions could be started by the injection of 4 vols mixture 1 onto 1 vol mixture 2. For experiments that involved MTX, PCPs were incubated in the presence of 5.57 mM DTT and in the presence or absence of 524 µM MTX and 524 µM NADPH (15 min at 21°C). Urea was added for a final concentration of 3.5 M, 10 minutes before addition to the import mixture (as 4 µl at 1.25 µM). Final concentrations of MTX and NADPH were 5 µM. For measurement of pep86 binding to 11S in solution, mitochondria were first solubilised by incubation with digitonin (5 mg/ml) at 4°C for 15 min. In selected experiments, depletion of Δψ was achieved by pre-treating mitochondria for 5 min with 10 nM valinomycin, and depletion of ATP was achieved by omitting ATP, creatine kinase and phosphocreatine from the reaction. ATP depletion was verified by monitoring sensitivity of mitochondria to a 5 min pre-treatment with 0.5 µM Antimycin A. PCP import is affected by Antimycin A when ATP is depleted but not under standard conditions. Luminescence was read from 125 µl reactions in a white round-bottom 96 well plate (Thermo Scientific) on either a CLARIOStar Plus (BMG LABTECH), or a BioTek Synergy Neo2 plate reader (BioTek Instruments) without emission filters. Measurements were taken every 6 seconds or less, and acquisition time was either 0.1 seconds (on the CLARIOStar Plus reader) or 0.2 seconds (on the Synergy Neo2 reader).

### Estimation of mitochondrial matrix volume

The mitochondrial matrix volume as a fraction of reaction volume was estimated using the previously published yeast mitochondrial matrix volume of 1.62±0.3 µl/mg (Koshkin and Greenberg, 2002). Thus when mitochondria are at 50 µg/ml, matrix volume will be 81±15 nl/ml, or ∼1/12345.68 total volume (between 1/15151.5 and 1/10416.7 accounting for error).

### Data processing and analysis

NanoLuc assay data were processed using a combination of software: Microsoft Excel, pro Fit 7 and GraphPad Prism versions 8 and 9. Data were then normalised to the maximum luminescence measurement for each experiment.

In most cases, the resulting data were fitted using pro Fit to a model for two consecutive, irreversible steps, where the final one gives rise to a signal (Fersht, 1984):

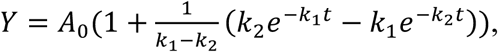

where A_0_ is amplitude, *k*_1_ and *k*_2_ the two rate constants, Y the signal and t is time. Note that this equation produces the same result whichever order *k*_1_ and *k*_2_ are in. Subsequent analyses of the resultant data were done in GraphPad Prism; linear and non-linear (Michaelis-Menten) regression.

### Membrane potential measurements with isolated mitochondria

Isolated mitochondria were diluted to 50 µg/ml in import buffer (described above) supplemented with 1 mM ATP, 0.1 mg/ml creatine kinase, 5 mM phosphocreatine, 10 µM GST-Dark protein and 0.5 µM Tetramethylrhodamine methyl ester (TMRM). Relative Δψ was monitored over time as a change in fluorescence of the Δψ-dependent dye TMRM in quenching mode. Fluorescence was measured at an excitation wavelength of 548 nm and an emission wavelength of 574 nm, in black plates, on a BioTek Synergy Neo2 plate reader (BioTek Instruments). The inner membrane PMF was generated by injecting 2 mM NADH, and PCP proteins added manually after stabilisation of fluorescence. Depolarisation was confirmed at the end of the assay by injecting CCCP.

## Acknowledgments

We would like to thank Prof. Andrew Halestrap for his insight and enthusiastic discussions on the mysteries of mitochondrial bioenergetics. We also thank past and present members of the Collinson lab who helped to get this project off the ground, particularly Drs Andrew Richardson and Dylan Noone.

## Author contribution

Project conceptualisation: GCP, HCF and IC

Sample preparation: HCF and XL

Data Collection: HCF

Data Analysis: HCF and WJA

Data interpretation: HCF, WJA and MSD

Manuscript writing: HCF, WJA, GCP and IC

Funding acquisition and project management: IC

## Declarations

The authors declare no competing interests. The funding agency and the University had no role in study design, data collection and analysis, decision to publish, or preparation of the manuscript.

For the purpose of Open Access, the author has applied a CC BY public copyright licence to any Author Accepted Manuscript version arising from this submission.

## FIGURES

**Figure 1 – figure supplement 1:**
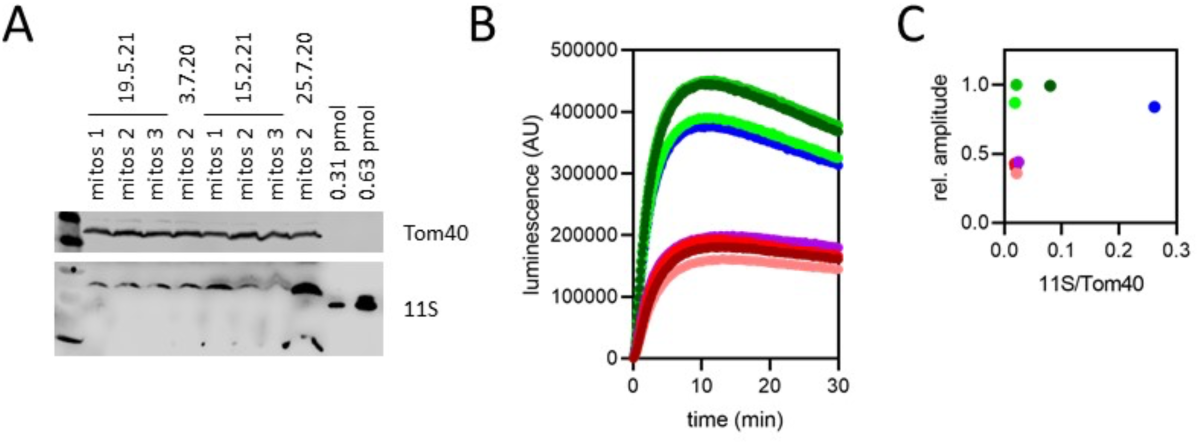
11S levels and signal amplitude. **A)** Western blot against 11S (bottom) and TOM40 (contro1, top) of eight different mitochondrial preparations extracted from four different batches of yeast. 60 µg each sample of mitochondria was fractionated by SDS-PAGE prior to Western blot. Two known concentrations of purified his-tagged 11S are also included for quantification by densitometry. Matrix concentration of 11S was calculated using the previously published yeast mitochondrial matrix volume of 1.62±0.3 µl/mg (Koshkin and Greenberg, 2002). **B)** Import traces of Acp1-pep86 with each of the mitochondrial preps in panel **A**, performed in parallel and unnormalised. **C)** Signal amplitude from panel **B** as a function of 11S concentration (normalised to TOM40) from panel **A**, with points coloured as in panel **B**. The results show no correlation between 11S concentration and amplitude.

**Figure 1 – figure supplement 2:**
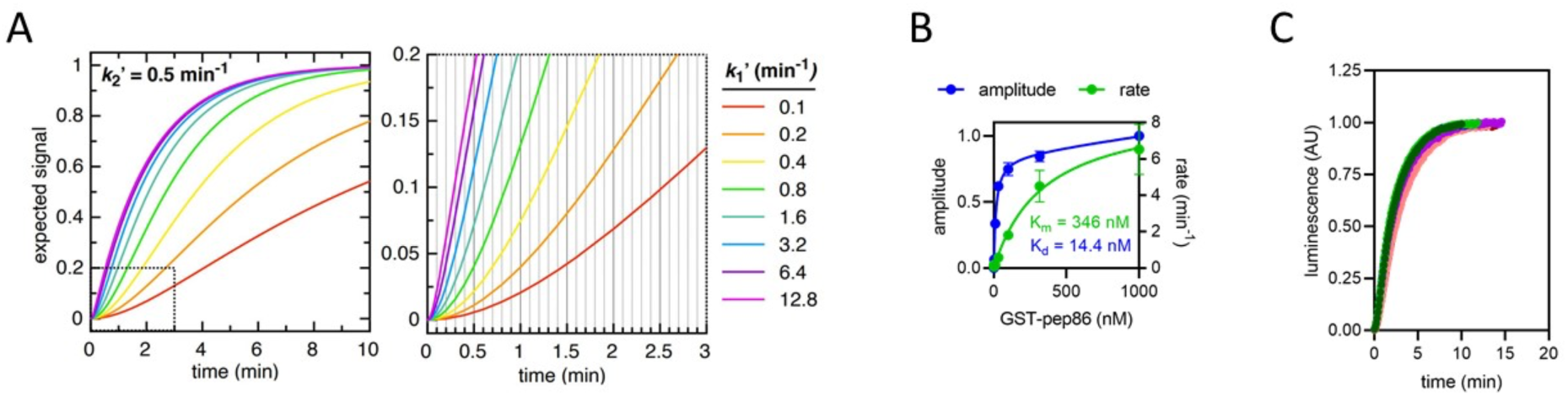
Constraints of data fitting to the NanoLuc import traces. **A)** The expected signal for a two-step import process, with *k*_2_′ fixed at 0.5 min^-1^ (for illustrative purposes) and *k*_1_′ varied between 0.1 min^-1^ (red) and 12.8 min^-1^ (magenta). As *k*_1_′ increases, it makes increasingly less difference to the overall shape of the curve. Because the plate reader measures luminescence with a frequency of 10 min^-1^ (represented as vertical gridlines in the zoomed in panel, right), any rate constants faster than about 5 min^-1^ will not be resolved. The same effect holds true for any additional rates that form part of the mechanism but are faster than ∼5 min^-1^. **B)** Amplitude (blue) and rate (green) determined from a single exponential fits to NanoLuc formation is solution. The pep86 tag is provided in the form of GST-pep86 which is not a PCP, and 11S comes from mitochondria solubilised completely with digitonin (5 mg/ml) to simulate binding within the mitochondrial matrix. Fits are to the Michaelis Menten equation giving an affinity of 14.4 nM and a *v*_max_ of 8.9 min^-1^. Data is shown as mean±SD of two independent biological experiments. **C)** The import traces in Figure 1 – figure supplement 1B all normalised to 1, coloured in the same way. For each trace, data collected at times after the maximum luminescence was recorded were excluded. The fact that all the traces overlay well confirms that binding of 11S is too fast to constitute either of the rates extracted from the two step fits – as expected given that the binding rate should be close to *v*_max_ for NanoLuc formation (as determined in panel **B**).

**Figure 3 – figure supplement 1:**
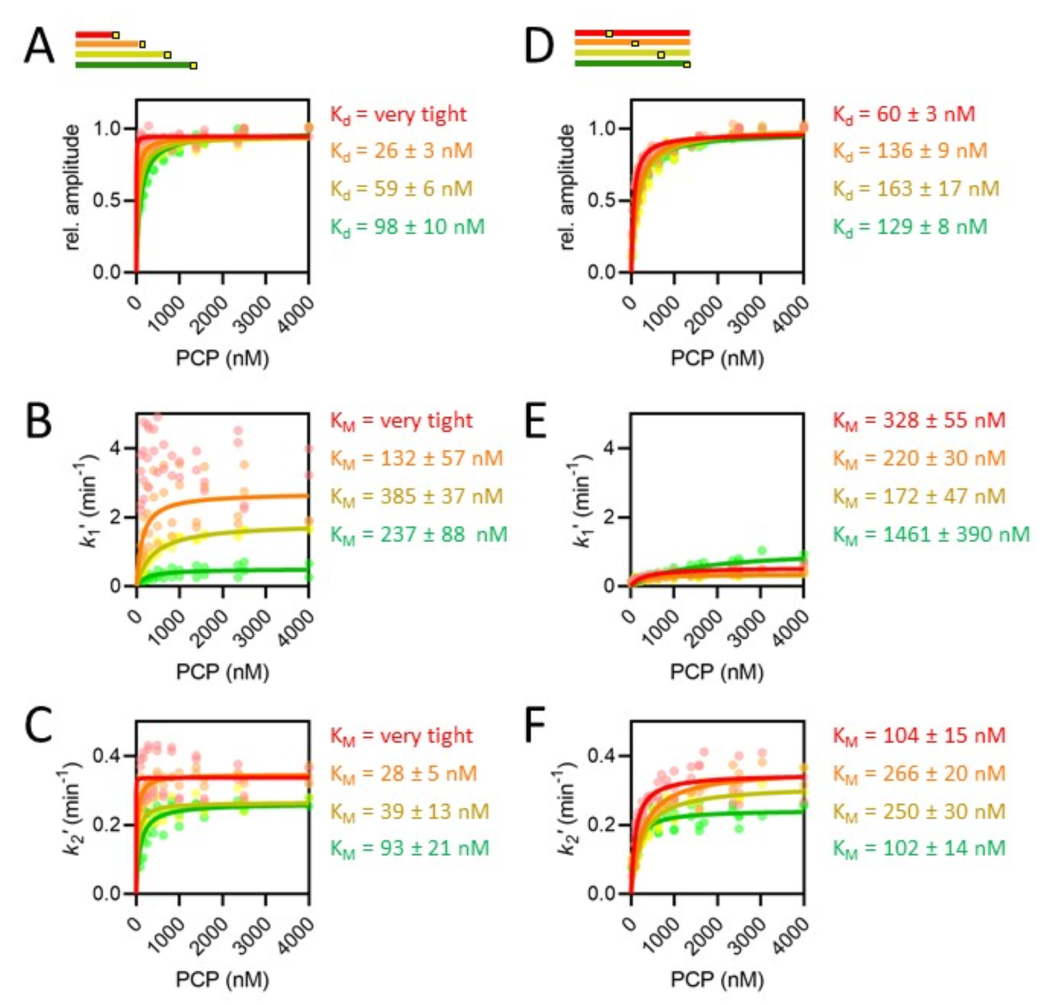
The concentration dependence of length and position variants. **A-F**) Amplitudes (**A, D**), *k*_1_′, assigned as the faster rate (**B, E**) and *k*_2_′ (**C, F**) for the length (**A-C**) and position (**D-F**) series, coloured red, orange, yellow and green in order of increasing length or pep86 position. All individual fits from 4-6 independent biological replicates of each set are shown, and the secondary data are fitted to the Michaelis-Menten equation, with errors estimated from the fitting.

**Figure 4 – figure supplement 1:**
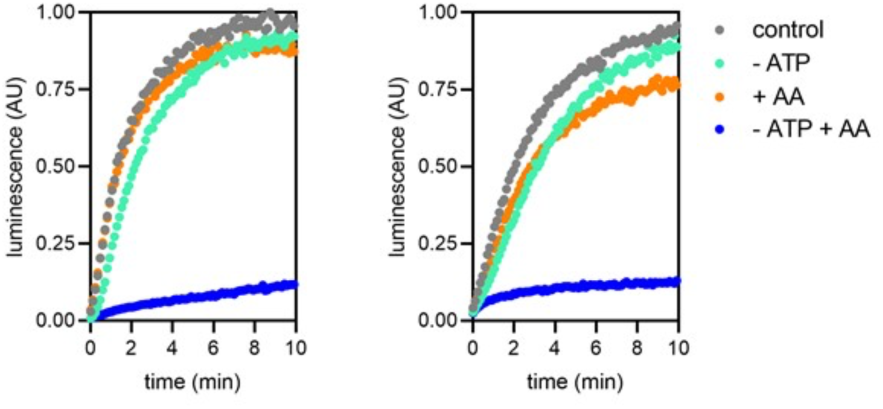
Confirmation of ATP depletion in the mitochondrial matrix. Import traces for 1 µM Acp1-pep86 (left) and Mrp21-pep86 (right) in the presence (grey and orange symbols) or absence (turquoise and blue symbols) of ATP and its regenerating system, and the absence (grey and turquoise) or presence (orange and blue) of antimycin A (AA).

**Figure 5 – figure supplement 1:**
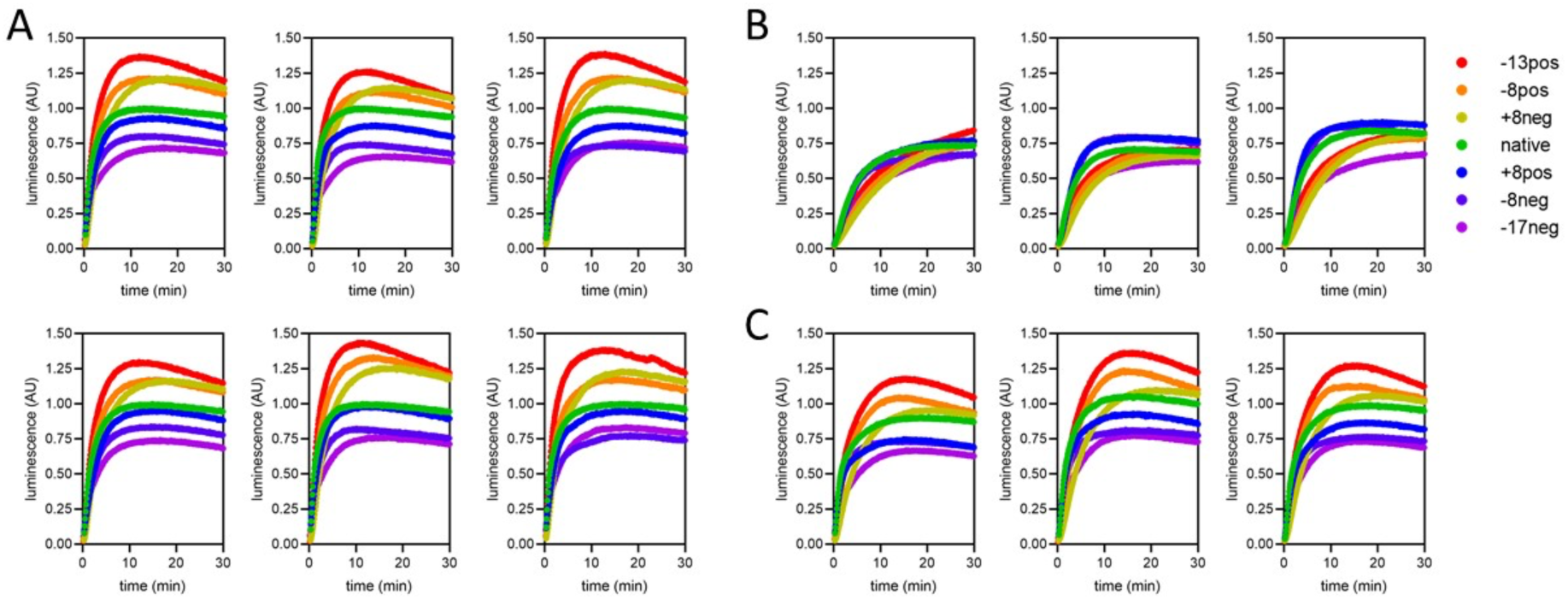
Complete import traces for the data in Figure 5. **A)** Two technical repeats each of three biological replicates, under standard conditions (1 µM PCP, ATP and regenerating system present and valinomycin absent). **B)** Three biological replicates with Δψ depletion (achieved by 5 min pre-treatment of mitochondria with 10 nM valinomycin). **C)** Three biological replicates with ATP depletion (achieved by excluding ATP and its regenerating system from the assay buffer)

**Supplementary Table 1.**
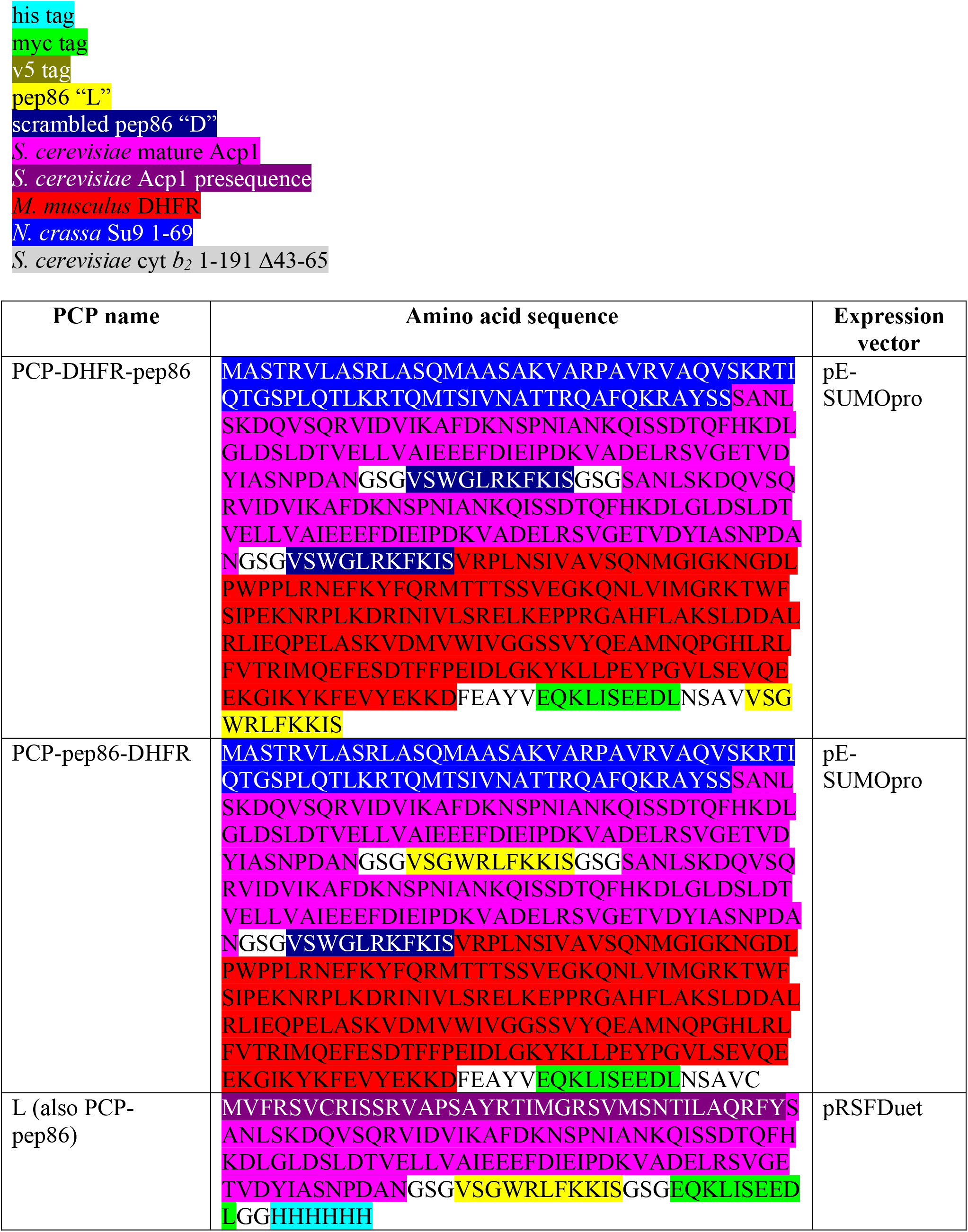
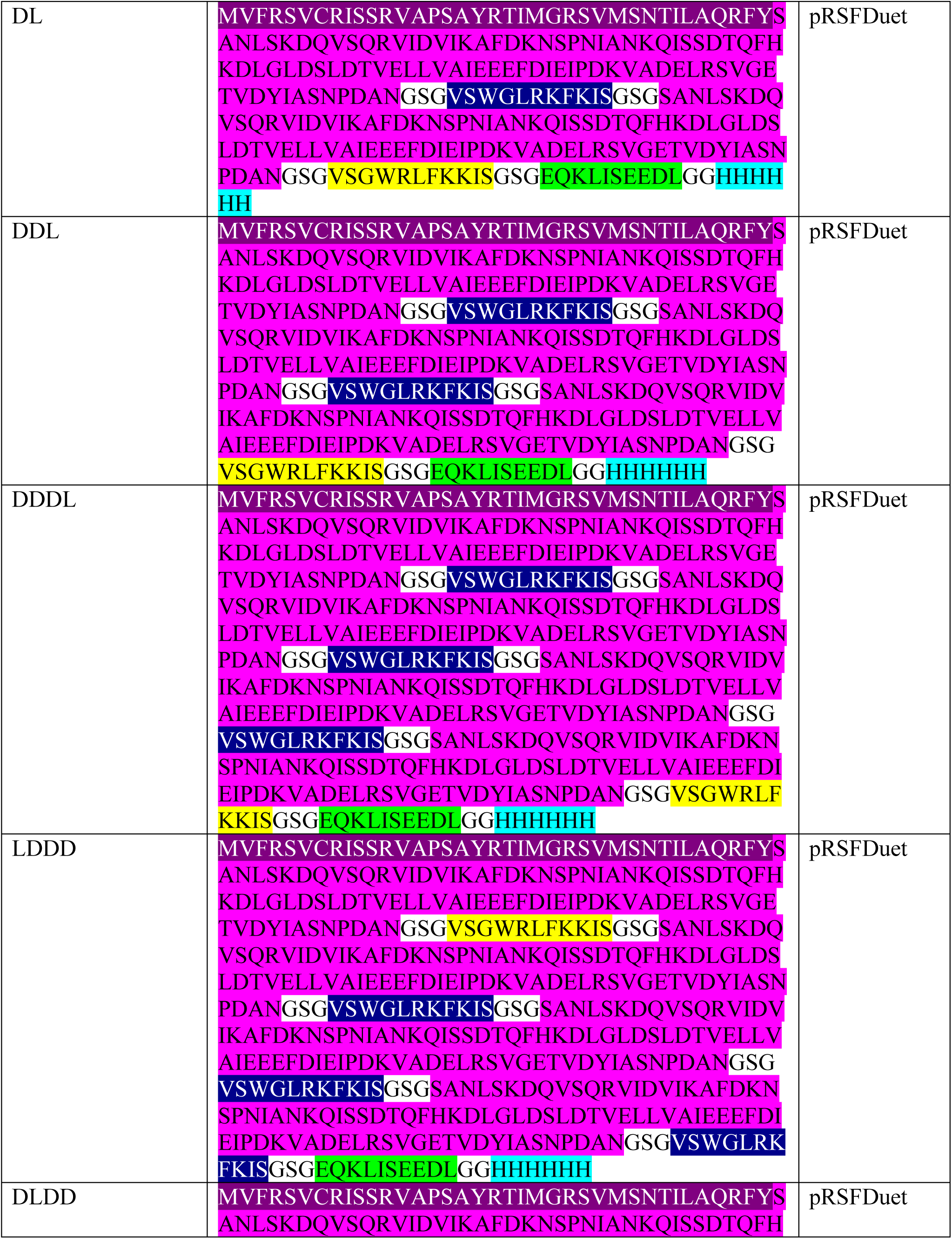
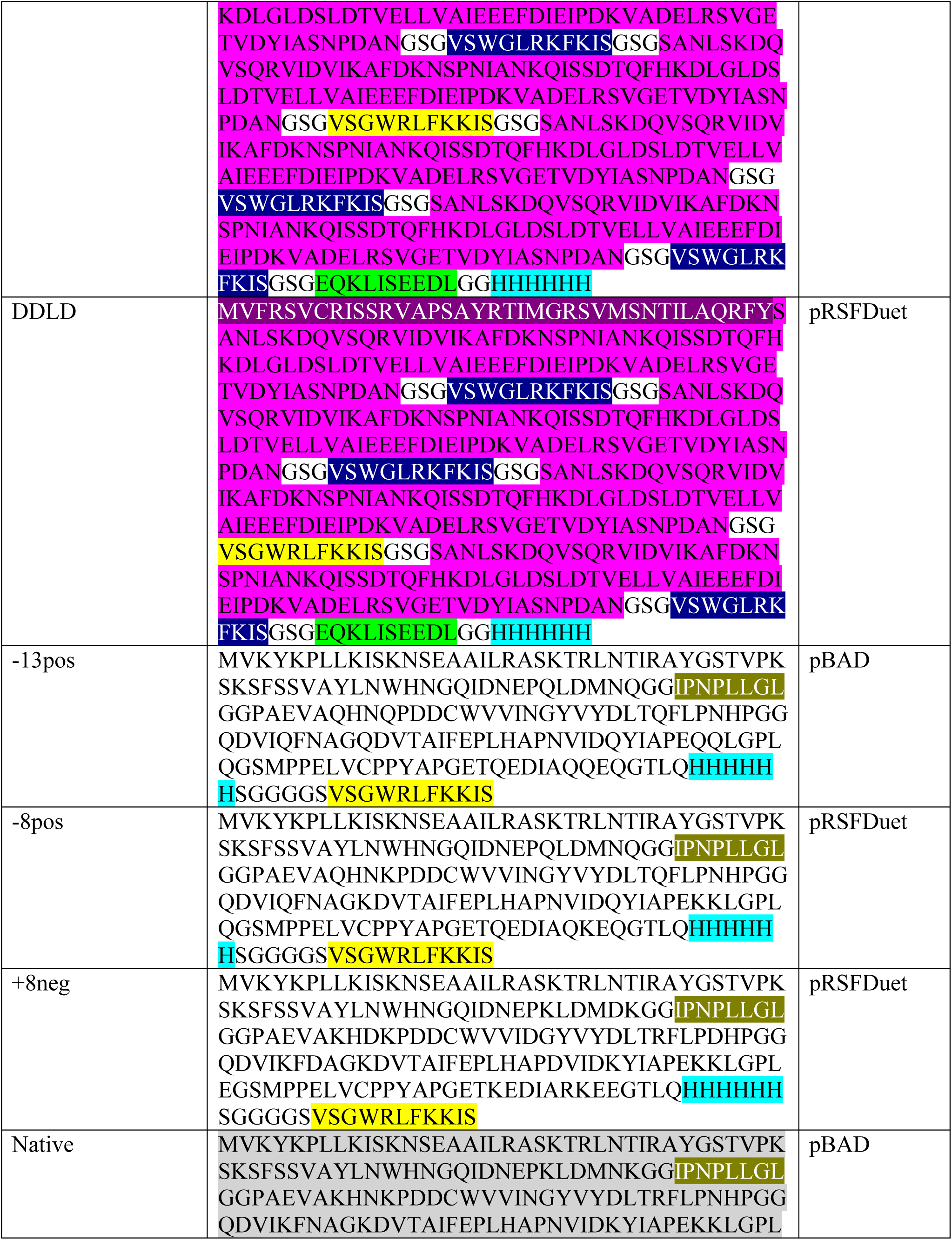
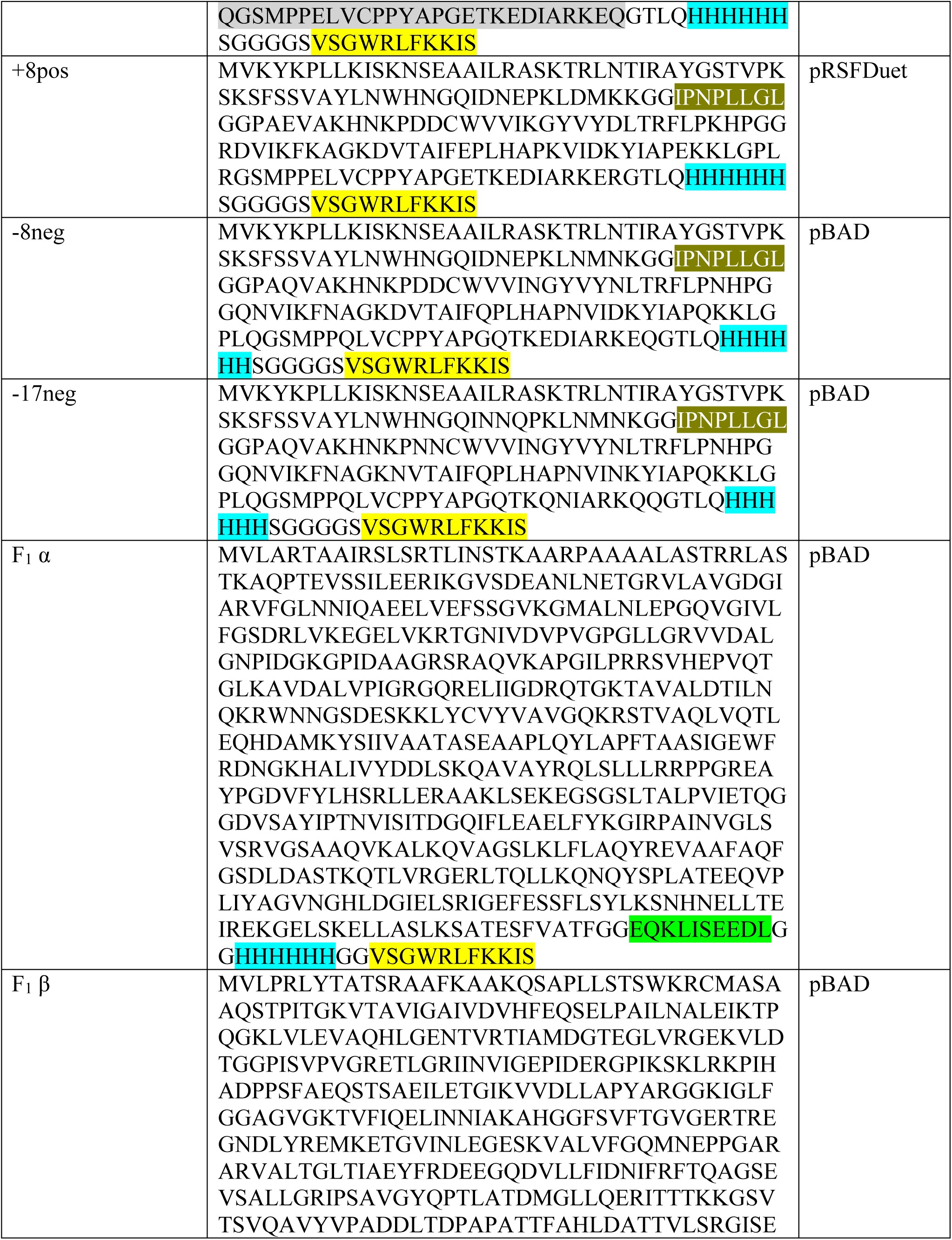
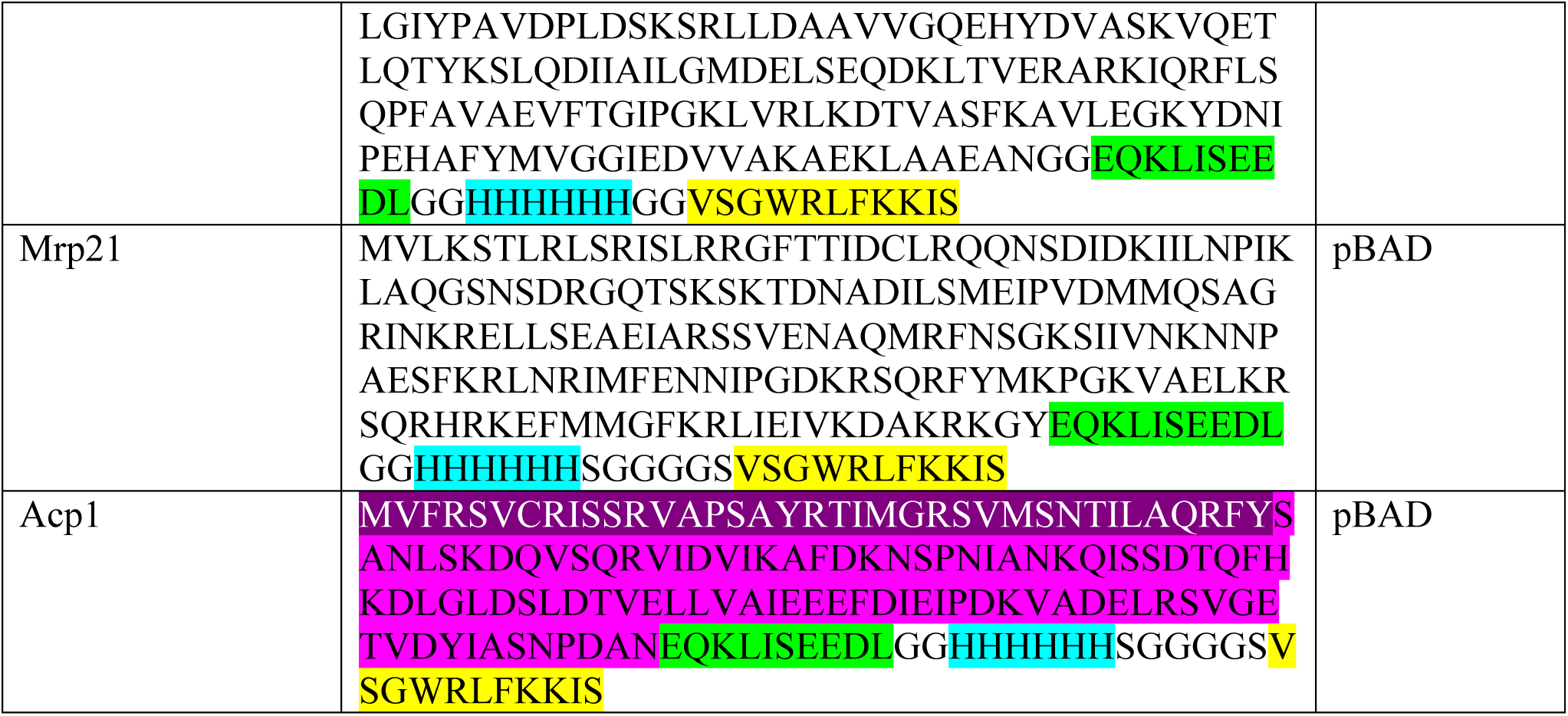

